# Naturalistic stimuli reveal a critical period in visual cortex development: Evidence from adult-onset blindness

**DOI:** 10.1101/2021.09.14.460305

**Authors:** Elizabeth Musz, Rita Loiotile, Janice Chen, Rhodri Cusack, Marina Bedny

## Abstract

How do life experiences impact cortical function? In people who are born blind, the “visual” cortices are recruited for nonvisual tasks such as Braille reading and sound localization (e.g., Collignon et al., 2011; Sadato et al., 1996). The mechanisms of this recruitment are not known. Do visual cortices have a latent capacity to respond to nonvisual information that is equal throughout the lifespan? Alternatively, is there a sensitive period of heightened plasticity that makes visual cortex repurposing possible during childhood? To gain insight into these questions, we leveraged naturalistic auditory stimuli to quantify and compare cross-modal responses congenitally blind (CB, n=22), adult-onset blind (vision loss >18 years-of-age, AB, n=14) and sighted (n=22) individuals. Participants listened to auditory excerpts from movies; a spoken narrative; and matched meaningless auditory stimuli (i.e., shuffled sentences, backwards speech) during fMRI scanning. These rich naturalistic stimuli made it possible to simultaneous engage a broad range of cognitive domains. We correlated the voxel-wise timecourses of different participants within each group. For all groups, all stimulus conditions induced synchrony in auditory cortex and for all groups only the narrative stimuli synchronized responses in higher-cognitive fronto-parietal and temporal regions. Inter-subject synchrony in visual cortices was high in the CB group for the movie and narrative stimuli but not for meaningless auditory controls. In contrast, visual cortex synchrony was equally low among AB and sighted blindfolded participants. Even many years of blindness in adulthood fail to enable responses to naturalistic auditory information in visual cortices of people who had sight as children. These findings suggest that cross-modal responses in visual cortex of people born blind reflect the plasticity of developing visual cortex during a sensitive period.

## Introduction

Studies with people who are blind provide key insights into how experience influences neuronal response properties. In congenitally blind individuals, the occipital lobes, which support visual processing in sighted people, are active during touch and audition (Collignon et al., 2011; Poirier et al., 2006; Sadato et al., 2002). “Cross-modal” responses are observed in a wide range of tasks, including perceptual tasks such as haptic shape recognition and motion detection (Gougoux et al., 2005; Uhl et al., 1991; Wanet-Defalque et al., 1988; Weeks et al., 2000). A number of studies have also documented responses during higher-cognitive tasks, including Braille reading, solving math equations and auditory sentence comprehension (Amalric et al., 2018; Bedny et al., 2011; Kanjlia et al., 2019; Lane et al., 2015; Röder et al., 2002; Sadato et al., 1996). Disrupting activity to “visual” regions impairs performance on Braille reading and verb generation, suggesting functional relevance of occipital cortex to nonvisual behavior in this population (Amedi et al., 2004; Cohen et al., 1999). In people born blind, responses to different cognitive processes are spatially dissociable within visual cortices and sensitive to subtle manipulations of higher-cognitive information, such as grammatical complexity of sentences and equation difficulty (e.g., Kanjlia et al., 2016).

Precisely how blindness leads to the emergence of these “cross-modal” responses in visual cortex remains unclear. One central question is whether cross-modal activity observed in congenital blindness is a consequence of functional cortical changes that are restricted to a sensitive period during development. According to one view, visual cortex has special potential to acquire nonvisual functions during time-sensitive windows early in life. Alternatively, nonvisual responses observed in people who are born blind could reflect latent capabilities of visual cortex that exist regardless of developmental experience and can be unmasked by vision loss at any age. Probing the response properties of visual cortices in individuals who became blind at different ages can provide data that distinguishes between these views. A particularly compelling source of evidence comes from studies with people who have lost their vision in adulthood. If a developmental sensitive period is a unique time during which visual cortices have the capacity to assume nonvisual functions, then people who lose vision as adults should not show the same cross-modal occipital recruitment as people born blind.

Currently available evidence relevant to this issue is mixed. On the one hand, there is evidence that responses to nonvisual tasks in the visual cortex are distinct in congenitally blind as compared to adult-onset blind people (for a review see Voss, 2013). While subregions of visual cortex are recruited during sound localization and motion processing in people who lost their vision in early childhood, such responses are reduced or absent in those who lost vision during adulthood (e.g., Burton, Snyder, Conturo, et al., 2002; Cohen et al., 1999; Collignon et al., 2013; Jiang et al., 2016; Sadato et al., 2002). Moreover, some evidence suggests that occipital responses are only behaviorally relevant in people who were born blind. TMS applied to the occipital lobes leads to increased error rates during a Braille reading task in congenitally blind but not adult-onset blind individuals (Cohen et al., 1999). Likewise, a number of studies have found decreased occipital recruitment during higher-cognitive tasks (i.e., math and language) in adult-onset blind individuals (Bedny et al., 2012; Kanjlia et al., 2019). There is also evidence that visual cortices of people who are born blind show subtle sensitivity to higher-cognitive information that is absent in people who become blind as adults. For example, only congenitally blind people show enhanced visual cortex activity for grammatically complex sentences over more simple sentence constructions and respond more for complex than simple math equations (Kanjlia et al., 2019; Pant et al., 2020).

At the same time, there is evidence that visual cortices of people who become blind as adults do respond to cross-modal tasks. Büchel and colleagues (1998) observed increased visual cortex activity during Braille reading in blind adults who lost their vision after puberty. In addition, in a series of studies, Burton and colleagues found that late-blind individuals activate occipital regions while participants listen to spoken words while making semantic judgments (Burton et al., 2002; Burton & McLaren, 2006) or generating verbs (Burton, Snyder, et al., 2002). Further, resting state analyses have revealed that both congenitally blind and adult-onset blind individuals show increased correlations between occipital cortices and fronto-temporal language areas (Butt et al., 2013; Sabbah et al., 2016), although these correlations are reduced in individuals who lost their vision as adults, relative to people born blind (Kanjlia et al., 2019). There is also an emerging literature suggesting that some cross-modal responses in visual cortices are observed even in sighted people. Studies have found increased visual cortex activity while sighted individuals perform tasks on nonvisual input, such as tactile discrimination (Merabet et al., 2008; Sathian & Zangaladze, 2002), haptic exploration of 3-D shapes (James et al., 2002), and even spoken sentence comprehension (Seydell-Greenwald et al., 2020). In addition, multivariate activity patterns in early visual cortex can distinguish between different natural sound categories in sighted people (Vetter et al., 2014). Taken together, these findings further raise the possibility that blindness during early life experiences is not necessary for cross-modal responses to emerge in the occipital cortices. Thus, the role of early life experiences in shaping visual cortex function remains unclear.

One challenge in addressing the question of whether visual cortices are more able to assume nonvisual functions during a critical period is the possibility that the functions assumed in adulthood are different from those assumed in childhood. In other words, people who become blind as adults could show just as much or more activity in visual cortices but for different cognitive functions as compared to people who are congenitally blind. Experiment designs that focus on a single cognitive domain may thus miss observing cross-modal responses in either congenitally blind or adult-onset blind people.

Naturalistic stimuli provide one way to approach this issue, by quantify brain responses across a broad range of stimulus features and cognitive demands simultaneously. Naturalistic auditory stimuli, such as spoken narratives and sound excerpts from live-action movies, are intrinsically rich with information across cognitive levels, ranging from low-level perceptual properties (e.g., changes in pitch and volume) to higher-order semantic content (e.g., plot and character intent). These stimuli have been shown to simultaneously engage multiple levels of the neurocognitive hierarchy, including sensory systems and functional networks that are implicated in linguistic processing and narrative comprehension (e.g., the default mode network; cf. Hasson et al., 2015; Lerner et al., 2011). As such, naturalistic stimuli are ideal for testing a broad range of possible responses simultaneously across different populations. In the current study we used such stimuli to quantify and compare cross-modal responses across congenitally blind, adult-onset blind and blindfolded sighted people.

In sighted people, a host of previous fMRI studies have found that listening to the same naturalistic stimulus induces correlated brain activity across people in multiple functional networks, including low level auditory areas and high-level semantic networks (Lerner et al., 2011; Simony et al., 2016). In effect, each participant’s activity timecourse serves as a model for the other participants. Cross-subject temporal synchronization indicates that in a given brain area, responses are time-locked to certain characteristics that fluctuate over the course of the stimulus presentation. For example, each time a speaker uses a complicated grammatical phrase, activity rises and falls at the same time across listeners in brain areas associated with language processing. In contrast, the auditory cortices synchronize to fluctuations in acoustic properties, e.g., volume. By systematically varying the content embedded in temporally extended naturalistic stimuli, researchers can infer the stimulus features that drive brain responses across different cortical networks (Hasson et al., 2015; Honey et al., 2012). Moreover, inter-subject correlations (ISC) provide a measure of the extent to which the same brain region serves a similar cognitive role across individuals (Hasson, 2004; Hasson et al., 2008). In order for synchrony to emerge in a given cortical location, the same time-varying stimulus features must drive activity in a similar way across different people. Thus, inter-subject correlations can quantify the degree to which the function of a given brain area is systematic.

Two studies have recently used naturalistic stimuli to elicit synchrony in visual cortices of congenitally blind people. Using magnetoencephalography, Van Ackeren and colleagues (2018) found that in congenitally blind but not sighted people, inter-subject activity in foveal V1 increased while participants listened to brief passages from audiobooks, relative to unintelligible speech. In addition, we recently showed that while listening to naturalistic auditory stimuli during fMRI, the visual cortices are more synchronized across congenitally blind than blindfolded sighted individuals (Loiotile et al., 2019). Critically, visual cortex synchrony among congenitally blind adults only emerged for stimuli that included meaningful content (e.g., sound excerpts from popular live-action movies and a spoken narrative). These same stimuli also synchronized higher-cognitive networks in fronto-parietal and fronto-temporal cortices in both sighted and blind groups. In contrast, meaningless sounds (e.g., backwards speech) only synchronized low-level auditory cortices but did not induce such synchrony in the visual system of congenitally blind adults. This finding suggests that in people who are born blind, the visual cortices are sensitive to meaningful and cognitively complex information embedded in auditory narratives. Furthermore, the reliable inter-subject correlations indicate that this reorganization systematically localizes to similar occipital regions across congenitally blind people (Loiotile et al., 2019; Musz et al., 2021).

In the present study, we leveraged this paradigm to investigate whether naturalistic auditory stimuli induce synchronized activity in the visual cortices of individuals who lost their vision as adults. Arguably, such stimuli provide an excellent opportunity to find cross-modal responses, regardless of which cognitive dimensions visual cortex is sensitive to in either congenitally blind or adult-onset blind people. By sampling a wide range of stimulus properties embedded in these stimuli, we can characterize the degree and type of cortical plasticity.

We can outline several possible predicted patterns of response to these stimuli in adult-onset blind individuals, although this is not an exhaustive list. One possibility is that visual cortex synchrony in adult-onset blind individuals is qualitatively and quantitatively similar to the synchrony previously reported in congenitally blind people (Loiotile et al., 2019). In other words, adult-onset blind individuals could show high synchrony for meaningful auditory stimuli i.e., audio-movies and narratives, but not meaningless auditory stimuli i.e., backwards speech. This result would suggest that like in people who are born blind, visual cortices of people who become blind as adults respond to higher-cognitive, meaning-related properties of naturalistic auditory stimuli. It would not necessarily indicate that the *same* cognitive aspects of these stimuli cause synchrony across these groups, however. For example, one could imagine synchrony caused by responses to language in one group and meaningful environmental sounds in another. In fact, it is also possible that visual cortex synchrony is even more robust among adult-onset blind people for meaningful auditory stimuli than in congenitally blind adults, if visual experience prior to adulthood facilitates visual imagery and this adds to the cross-modal responses in visual cortex. Relative to other types of auditory stimuli, naturalistic meaning stimuli, such as auditory movies, are arguably most likely to elicit such imagery. Such a pattern would suggest that cross-modal responses in visual cortex of blind individuals do not require blindness during development but instead can be unmasked at any age by lack of visual input to occipital cortex.

An alternative possibility is that visual cortex synchrony in adult-onset blind individuals is equally robust but cognitively distinct from that observed in congenitally blind people. For example, we might find that synchrony in adult-onset blind individuals is less selective, and observed not only for meaningful auditory stimuli, such as audio movies, but also for meaningless auditory stimuli, including backwards speech. This would suggest that cross-modal recruitment in visual cortex is possible throughout the lifespan but recruitment in adulthood is related to sensory auditory processing, rather than higher-cognitive processing.

Finally, it is possible that, even for rich naturalistic stimuli, visual cortex responses in adult-onset blind people are reduced relative to people who are congenitally blind and no different from those who are sighted. Such a result would suggest that there is a sensitive period of development during which the visual cortices are uniquely able to enhance their responses to nonvisual information. Cross-modal responses in people who lose vision as adults are thus comparatively weak or inconsistent across individuals, relative to those who lost their vision in infancy. It is important to note that in our previous study with the same naturalistic stimuli, we found that even sighted individuals show some degree of synchrony in visual cortices. This synchrony is weak, relative to the congenitally blind group, and not equally selective. Nevertheless, synchrony observed in blindfolded sighted individuals suggests that some nonvisual information reaches the occipital cortices in all people regardless of vision status. The goal of the present study was therefore to ask whether the pattern of synchrony observed in adult-onset blind individuals is more like that of congenitally blind people or like that of people who are sighted. In other words, does long-term lack of vision in adulthood significantly boost cross-modal recruitment beyond what can be achieved by approximately an hour of blindfolding during the scan?

In order to test among these possibilities, we measured brain responses while participants from three vision groups (blindfolded sighted, congenitally blind, and adult-onset blind) listened to naturalistic auditory stimuli, including 5-7 minute long excerpts from three popular live-action movies and a spoken narrative. Participants also listened to semantically degraded versions of the spoke narrative, one in which the linguistic content was preserved by the coherent plot line was removed (by shuffling the individual sentences in the narrative), and a version that lacked linguistic content but retained acoustic features (by playing the narrative in time-reverse format). We then tested for synchronized activity across vision groups and across stimulus conditions, both in visual cortex and throughout the brain.

## Methods

### Participants

Twenty-two sighted (S, 4 males), fourteen adult-onset blind (AB, 5 males) and twenty-two congentially blind (CB, 8 males) participants contributed data to this study (Table 1). All participants were blindfolded during testing. All blind participants reported having at most minimal light perception at the time of the experiment, and had vision loss due to pathology at or anterior to the optic nerve, and not due to brain damage (Table 2). Sighted participants reported normal or corrected-to-normal vision. Participants with adult-onset blindness became blind after the age of 19 (mean = 38.1, SD = 13.9, min = 19, max = 57) and were blind for an average of eighteen years after reaching their current level of vision loss (SD = 10.8, min= 6, max= 40) (see Table A.1 for demographic information). Average age of participants in the AB group was greater than both the CB, *t*(34)= 2.33, *p*= .03 (AB: mean= 54.3 years, *SD*= 9.7; CB: mean= 43.5, *SD*= 16.9), and S groups *t*(34)= 2.86, *p*= .01 (S: mean= 42.7, *SD*= 13.8), which were matched to each other. Data from the sighted participants and congentially blind participants were previously published (Loiotile et al., 2019; Musz et al., 2021).

**Table 1.**
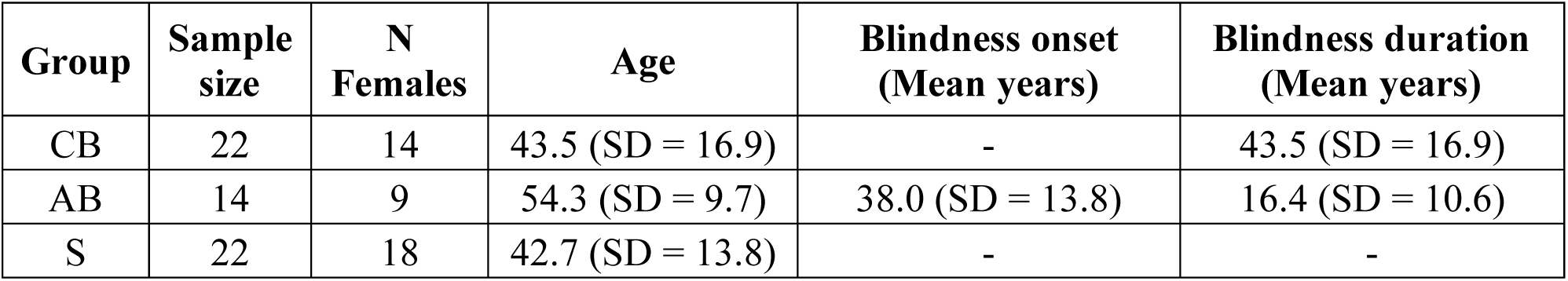
Participant demographic information and vision loss history summary for the congenitally blind (CB), adult-onset blind (AB) and sighted (S) groups. Duration of blindness is calculated by subtracting age at testing from age at which the current level of vision was reached for the AB group, and age at time tested for the CB group.

**Table 2.** Etiology summary for the congentially blind (CB) and adult-onset blind (AB) groups showing causes of blindness.

| Group | Blindness etiology | Participant number |
| --- | --- | --- |
| **CB** | Retinopathy of Prematurity | 8 |
|  | Leber Congenital Amaurosis | 7 |
|  | Optic nerve hypoplasia | 3 |
|  | Retinitis Pigmentosa | 1 |
|  | Unknown | 3 |
| **AB** | Glaucoma | 3 |
|  | Retinitis Pigmentosa | 3 |
|  | Diabetic retinopathy | 2 |
|  | Trauma | 2 |
|  | Optic nerve neuropathy | 1 |
|  | Stevens-Johnson syndrome | 1 |
|  | Uveitis | 1 |
|  | Autoimmune disorder | 1 |

### Stimuli

Participants listened to six audio clips, each five to six minutes in duration. Three of the audio clips were excerpted from popular live-action movies (*Blow-Out, The Conjuring*, and *Taken*). These Audio-Movie clips were chosen because they are engaging, suspenseful, and easy to follow, and could therefore facilitate a shared interpretation and experience across particpants. The three other audio clips were excerpted from the same six-minute excerpt of a live comedy sketch (“Pie-Man”). The intact version was presented in its original format. The “Sentence-Shuffle” version was created by splicing and re-ordering the sentences from the intact version, thereby retaining intelligible sentences but removing the coherent plotline. The “Backwards Speech” version was created by playing the excerpt in time-reverse format, such that it lacked any intelligble content. In addition, subjects also participated in a resting state scan, during which no stimuli were presented and participants were instructed to relax and stay awake. All stimuli are available for download at the Open Science Framework: https://osf.io/r4tgb/

### Procedure

During fMRI scanning, participants listened to each sound clip, with one clip presented during each of six scanning runs. Prior to scanning, we confirmed that each participant had not previously seen the movies that the clips were sourced from. Clips were presented in pseudo-random order, where each presentation sequence for an AB participant was yoked to a corresponding S participant and CB participant. Participants were instructed to pay attention and listen carefully, as they would have to answer questions about the plot details of each sound clip at the end of the experiment. Immediately before each scanning run, participants were read a brief prologue that provided context for the upcoming sound clip. The presentation of each sound clip was preceded by five seconds of rest and followed by 20-22 seconds of rest. These rest periods were excluded from all analyses. Due to time constraints, one sighted participant and four AB participants did not complete the resting state scan.

Following fMRI scanning, participants immediately received an orally-presented post-scan questionnaire composed of five multiple choice questions for the comedy sketch and each movie clip (see Table A2 for behavioral performance by subject group). We included data from all participants in all reported brain analyses. While the mean scores for the AB group were consistently lower than the scores for the CB and S groups, post-hoc t-tests revealed that these differences were not statistically significant (all *p*s>.05). In order to most closely match the sample size and behavioral performance of the AB (n=14) and CB (n=22) groups, we repeated all brain analyses in a subset of CB participants (n=14) whose behavioral performance mostly closely matched the AB group (Table A2). Brain results for the subset CB group are qualitatively similar to the results that include all CB participants (Figure 1).

**Figure 1.**
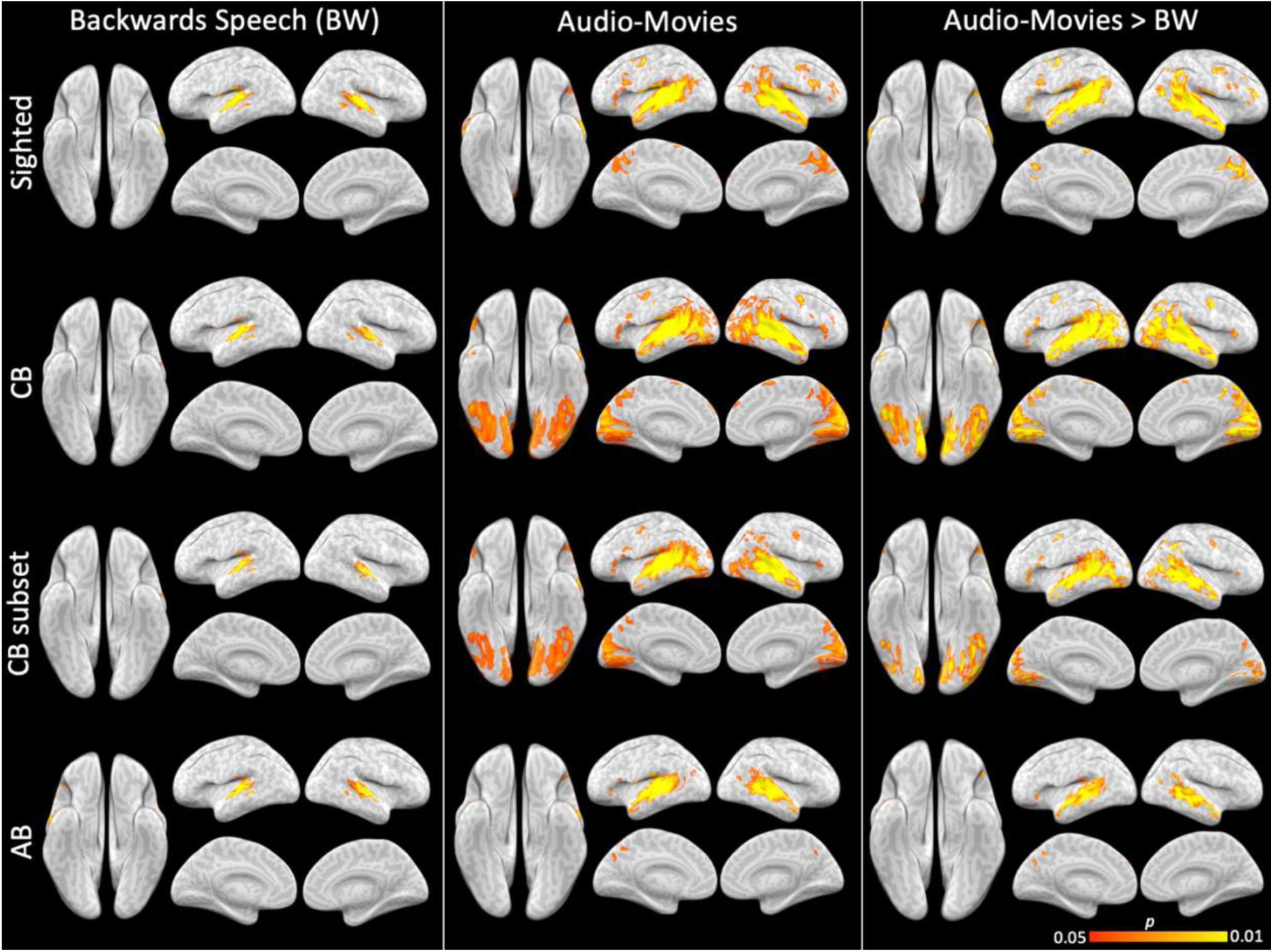
Whole-brain inter-subject correlation (ISC) reliability maps for the Backwards Speech stimulus (BW) and the Audio-Movie stimulus in each subject group. Voxel-level synchronization is shown within the sighted (S) group; the congenitally blind (CB) group; a subset of the CB group (CB subset) that matches the size of the adult-onset blind (AB) group (both n=14); and the AB group. Maps showing above-threshold ISC for each individual stimulus condition (first and second columns) are corrected for multiple comparisons. For each subject group, the corresponding difference map (third column) is limited to voxels that exceeded the threshold for either individual stimulus condition.

Auditory stimuli were presented over Sensimetrics MRI-compatible earphones (http://www.sens.com/products/model-s14/) at the maximum comfortable volume for each participant. To ensure that participants could hear the softer sounds in the auditory clips over the scanner noise, a relatively subtle, low-frequency sound (RMS amplitude = 0.002, frequency = 3479) was played to participants during acquisition of the anatomical image; all participants indicated hearing the sound via button press. Participants were told to press a button at any point during the scanning session if they could not hear the contents of the sound clips over the noise of the scanner. In such cases, we ended the scanning run, re-adjusted the audio volume, and performed additional sound checks. The scanning run and corresponding sound clip were then restarted.

### fMRI data acquistion

Whole-brain structural and functional MRI data were collected on a 3 Tesla Phillips scanner. T1-weighted structural images were collected in 150 axial slices with 1 mm isotropic voxels using a magnetisation-prepared rapid gradient-echo (MPRAGE). T2*-weighted functional images were collected with a gradient echo planar imaging sequences in 36 sequential ascending axial slices with 2.4 × 2.4 × 3 mm voxels and 2-second TR (echo time: 3ms, flip angle: 70 degrees, field-of-view: 76×70 matrix, slice thickness: 2.5mm, inter-slice gap: 0.5, slice-coverage: FH 107.5, PH direction L/R, first-order shimming).

### fMRI data processing

Preprocessing was performed using FEAT (fMRI Expert Analysis Tool) Version 6.00, part of FSL (http://fsl.fmrib.ox.ac.uk/fsl) and included slice time correction using Fourier-space time-series phase-shifting, motion correction using MCFLIRT (Jenkinson et al., 2002), high-pass filtering (140s), and linear detrending. All functional volumes were co-registered and affine transformed to a standard anatomical brain (MNI152) using FLIRT. Functional images were smoothed with a 4mm FWHM Gaussian kernel and resampled to 3mm isotropic voxels. The first four and last eight TRs of each scanning run were discarded, corresponding to the rest periods before and after the clips (accounting for delays in hemodynamic response). Analyses were performed in volume space and data were projected onto a cortical surface with NeuroElf for visualization (neuroelf.net).

### Inter-subject correlation analysis

We used inter-subject correlation (ISC) to test whether the same anatomical locations perform consistent functions across individuals. For each vision group and each auditory stimulus, we assessed the degree of stimulus-driven synchonization across individuals at each voxel in the brain. ISC is defined as the correlation of BOLD activity timecourses across participants as they listen to or view a common stimulus (Hasson et al., 2004; Hasson et al., 2010). Within-group ISC (i.e., “reliability”) was calculated as the average correlation at each voxel between that voxel’s BOLD activity timecourse in one individual subject and the average of BOLD timecourse in the remaining individuals in the group. Between-group ISC was calculated as the correlation between the timecourse from each subject in one group (e.g., a participant AB group) versus group-average timecourse of the comparison group (e.g., the mean timecourse of the CB group). Subject versus rest-of-group correlations (or subject versus comparison group correlations, for the between-group comparisons) were then averaged together, resulting in one mean ISC value at each voxel. The voxel-level ISC values were then projected to the brain to yield group-level ISC maps. To create mean Audio-Movies synchronization maps, the averaged *r*-value ISC maps for each movie condition were first transformed to Fisher’s *z*-values. The three resulting *z*-maps (one per movie) were then averaged together and subsequently transformed back into *r*-maps.

To evaluate synchrony among subjects for each stimulus condition, the statistical likelihood of each observed ISC value was assessed using a bootstrapping procedure based on phase-randomization, and maps were corrected for multiple comparisons using non-parametric family-wise error rate (Regev et al., 2013). All ISC and statistical analyses were performed with custom software written in MATLAB (MathWorks). The null hypothesis was that the BOLD response timecourse at each voxel in each individual was independent of the BOLD response timecourses in the corresponding voxel of the other individuals (i.e., that there was no inter-subject reliability among individuals). For each scanning run and stimulus condition, we applied a phase randomization to each voxel timecourse by applying a fast Fourier transform to the signal, randomizing the phase of each Fourier component, and then inverting the Fourier transformation (Lerner et al., 2011; Loiotile et al., 2019). This procedure scrambles the phase of the timecourse but preserves its power spectrum. For each randomly phase-scrambled surrogate dataset, we computed ISC for all voxels in the same manner as for the empirical correlation maps described above. The resulting correlation values were then averaged across all subjects within each voxel, yielding a null distribution of mean ISC values for all voxels.

To correct for multiple comparisons, we retained the highest ISC value from the null distribution of all voxels in a given iteration. This bootstrapping procedure was repeated 1000 times to obtain a null distribution of maximum noise correlation values. Familywise error rate was defined as the top 5% of the null distribution of the maximum correlation values, which was then used to threshold each veridical map (Nichols & Holmes, 2002). Thus, we rejected the null hypothesis for a particular comparison if an observed ISC value was in the top 5% of all 1000 values in each null distribution (i.e., a one-tailed statistical test). Differences in thresholds across stimulus conditions and subject groups (values ranging from 0.09 to 0.15) reflect variances in the null sampling distributions, due to differences in the degrees of freedom among stimuli (e.g., number of timepoints) and between the sample sizes across groups. To conservatively test whether the visual cortices in the AB group show any synchronization in response to nonvisual stimuli, for each stimulus condition we applied the most lenient threshold from the three subject groups. Results were qualitatively similar to those obtained by using each group’s own criteria.

In order to identify voxels that show an increase in ISC for one stimulus condition over the other within each subject group (e.g., Audio-Movies versus Backwards Speech among CB individuals), a paired *t*-test (two-sided, alpha = 0.05) was performed at each voxel that was above-threshold in at least one of the two compared conditions. Thus, the *t*-test was performed by comparing each participant’s two ISC values from each condition at each included voxel.

### Inter-subject correlation: Region of interest (ROI) analysis

We used three bilateral ROIs: primary visual cortex (V1), the early auditory cortex (A1), and higher-cognitive posterior lateral temporal cortex (PLT) (Loiotile et al., 2019). The V1 ROI was sourced from a previously published anatomical surface-bsaed atlas (PALS-B12; Van Essen, 2005). The early auditory cortex ROI was defined as the transverse temporal portion of a gyral based atlas (Desikan et al., 2006; Morosan et al., 2001). A higher-cognitive bilateral PLT ROI was taken from parcels responding to higher-level lingustic conetent in sighted subjects, defined as responding more to sentences versus non-word lists (Fedorenko et al., 2010). This PLT subregion is sensitive to high-level lingusitic information, ranging from word and sentence-level meaning to sentence structure (Bookheimer, 2002).

For each ROI, ISC was computed across participants using each individual’s response timecourse, averaged across all voxels in the ROI. Statistical significance was assessed with the same procedure that was applied to the whole-brain analysis. Namely, *p*-values were computed by comparing each observed ISC value to a null distribution of noise correlation values that were created by applying phase randomization to each ROI timecourse. No subsequent multiple comparisons correction were performed. All statistics for factor comparisons (i.e., ROI, group, and/or conditions) were obtained by subtraction of the relevant *z*-transformed ISC values. Fisher’s *z*-transformed-*r* values were subsequently transformed back to *r* (correlation coefficient) values for reporting.

To test for group differences in ISC after accounting for variance attributed to participant age, regression analyses were run using participant age to predict participant-level ISC values in a given ROI. Then, residuals from the regression model were compared across groups. Group-average differences in these residuals were then computed. To assess statistical significance, these values were compared to a null distribution of group differences in such residual values, which were derived in the same manner but using the phase-randomized timecourses to compute ISC instead. Each regression model included data from from two subject groups at a time (i.e., AB vs. CB; AB vs. S) to enable pairwise comparisons.

To assess whether blindness duration among the AB group and the CB group contributes to a participant’s timecourse synchrony to other individuals in visual cortex, we correlated each blind participant’s duration of blindness with their individual ISC value for the Audio-Movies condition in the V1 ROI. For AB individuals, this correlation was computed two different ways: once using each participant’s synchrony to the rest-of-AB-group average timecourse (i.e., within-group ISC) and once using each participant’s synchrony to the CB group average timecourse (i.e., between-group ISC). For the CB group, the correlation was computed between blindness duration (i.e., age) and within-group ISC (i.e., each participant’s synchrony to the rest-of-CB group average timecourse).

## Results

### Audio-Movies and intact Spoken-Narrative induce synchrony in higher-cognitive areas in all subject groups, relative to Backwards Speech

#### Audio-Movies

In response to the Audio-Movies stimuli, significant inter-subject synchrony emerged in all three subject groups in several nonvisual brain areas associated with higher cognition. In particular, the Audio-Movie stimuli induced significant ISC across bilateral temporal cortex, including the temporal poles, superior and middle temporal gyrus and the angular gyrus. Synchrony also emerged in the lateral prefrontal cortex and on the medial surface in the precuneus (Figure 1). As previously reported, synchrony observed in these nonvisual regions was similar across the CB and S groups (Loiotile et al., 2019). The AB group showed reliable ISC in similar higher-cognitive regions, including left precuneus and bilateral lateral temoral cortex, although synchrony among AB individuals in these areas was less spatially extensive than in the other groups.

In contrast to the Audio-Movies, the synchrony induced by the Backwards Speech stimuli was limited to bilateral auditory cortex in all three groups (Figure 1). In direct comparisons between Audio-Movies versus Backwards Speech, ISC was greater for the Audio-Movies in each group in the same regions that showed above-threshold synchrony for the Audio-Movies condition, including bilateral prefrontal cortex, lateral temporal cortex, and precuneus. No brain area showed the reverse contrast, with greater ISC for Backwards Speech than Audio-Movies.

#### Spoken-Narrative

For the intact version of the spoken narrative (“Pie-Man”), the synchrony in nonvisual regions largely matched the results for the Audio-Movies. For all three subject groups, Pie-Man induced synchronized responses in bilateral temporal cortex. For the CB group and the S group, but not the AB group, synchrony also emerged in precuneus and lateral prefrontal cortex (Figure B1). In addition, both the CB and AB groups also showed small clusters of reliable ISC along the medial surface of the frontal cortex. This synchrony was bilateral in the CB group and right-lateralized in the AB group.

In contrast to the intact version of Pie-Man, synchrony for the Sentence-Shuffle stimulus condition was limited to the bilateral temporal cortex in each of the three subject groups (Figure B1). In both the CB and the S groups, synchrony in lateral temporal cortex did not extend posteriorly to the angular gyrus or anteriorly to the temporal pole, as was observed for the intact Pie-Man and the Audio-Movies. In contrast, the Sentence-Shuffle and Pie-Man ISC maps for the AB blind group were largely similar.

As expected, the resting state scan did not induce synchrony among individuals in any brain region in any subject group.

### Audio-Movies and intact Spoken-Narative stimuli synchronize responses in the visual cortices of congenitally blind, but not sighted or adult-onset blind individuals

#### Audio-Movies

Within the CB group, but not in the S group or AB group, the Audio-Movies synchronized activity in the bilateral occipital cortices. In the CB group exclusively, reliable ISC spanned the medial, lateral, and ventral surfaces of visual cortex (Figure 1). This pattern of results was recapitulated in the ISC maps yielded by each individual movie clip (Figure B2). Each audio-movie synchronized responses in visual cortices, but only among the CB participants. In contrast, the Backwards Speech stimulus did not induce synchrony in visual regions of any subject group. The direct contrast of Audio-Movies versus Backwards Speech ISC showed that synchrony in the visual cortices was reliably greater for the Audio-Movies, but only for the CB group.

Since there were more CB than AB participants, we performed a subset analysis to ensure that difference among groups was not introduced by the difference in sample size. For this analysis, we generated ISC maps using a subset of CB participants of equal size to the AB group (Figure 1, third row). Like the AB group, the CB subset map showed less spatially extensive synchrony in the higher-cognitive nonvisual regions for the Audio-Movies, relative to the S group and the full CB group. However, unlike the AB group, the CB subset group still showed visual cortex synchrony while listening to the Audio-Movies, and not during Backwards Speech. Thus, group size could not account for visual cortex synchrony differences across CB and AB groups.

#### Spoken-Narrative

For the CB group but not the S group or the AB blind group, the intact Pie-Man stimulus induced synchronized responses in the bilateral occipital cortices (Figure B1). The location and spatial extent of this synchrony in visual cortex largely matched the above-threshold ISC locations observed for the Audio-Movies. In contrast, the Sentence-Shuffle condition did not induce visual cortex synchrony for any of the three subject groups.

### ROI Analysis: Higher V1 synchrony in congenitally blind group, relative to adult-onset blind and sighted

To more closely examine how inter-subject synchrony varied across subject groups and stimulus conditions in V1, we conducted ROI analysis (Figure 2). As a point of comparison, we also examined synchrony in early auditory cortices (A1) and language-responsive posterior lateral temporal (PLT) cortex. Since there is a large number of possible comparisons between groups and stimuli, increasing a false-positive risk, we conducted only hypothesis driven comparisons.

**Figure 2.**
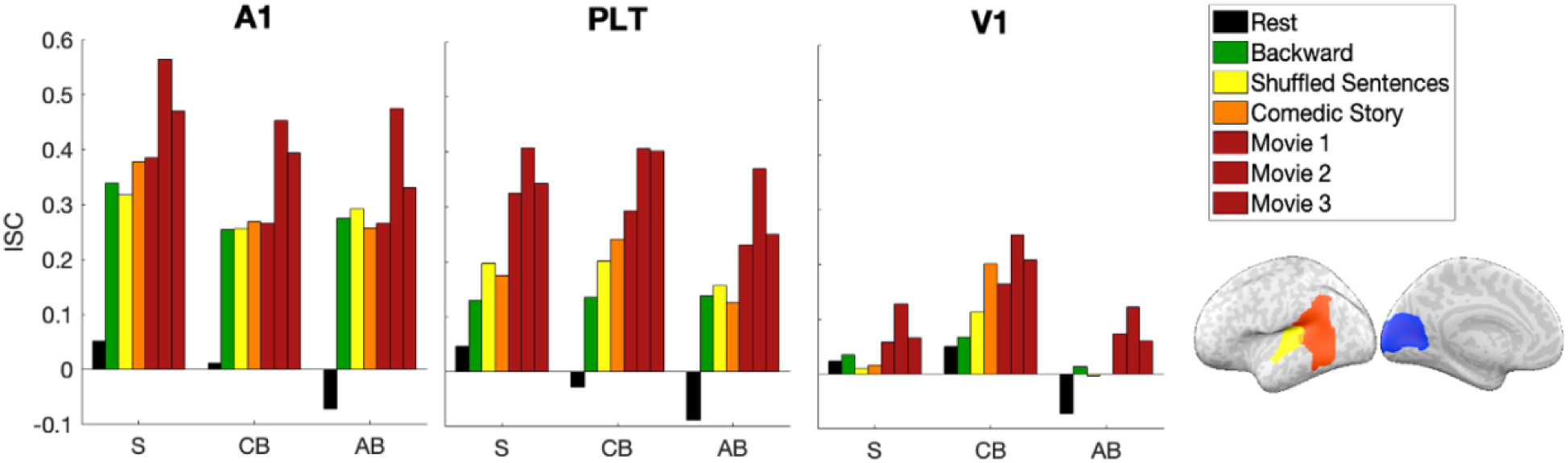
Left: Within-group ISC values for each region of interest (ROI) and each subject group. Right: ROIs are displayed in the left hemisphere (A1 in yellow; PLT in orange; V1 in blue) but inter-subject correlations are calculated bilaterally. Results for each individual audio-movie appear in the order listed and are shown for illustration purposes only; the reported results for Audio-Movies are averaged across the three movies. A1= early auditory cortex; PLT = posterior lateral temporal cortex; V1 = primary visual cortex

In V1, Audio-Movies elicited reliable ISC in all three groups (S: *r*=.08; CB: *r*= .21, AB: *r*=.08, all *p*>.001), while the Backwards Speech condition only showed above-chance synchrony in the CB group (*r*=.07, *p*=.01). Audio-Movies induced greater ISC than Backwards Speech in all three groups (all *p*<.001). Audio-Movies synchrony for the CB group was significantly greater than the S (*p*<.001) and the AB groups (*p*<.001), while the AB and S groups did not differ from each other (*p*>.1). In tests for group-by-condition interactions, the increase in ISC for Audio-Movies versus Backwards Speech was greater in the CB group than the S group (*p*<.01) and marginally greater in the CB group than the AB group (*p*=.07), while the difference between the AB and S groups was not significant (*p*>.1). Participant-level V1 ISC values in response to the Audio-Movie stimuli are shown in Figure 3 and Figure C1.

**Figure 3.**
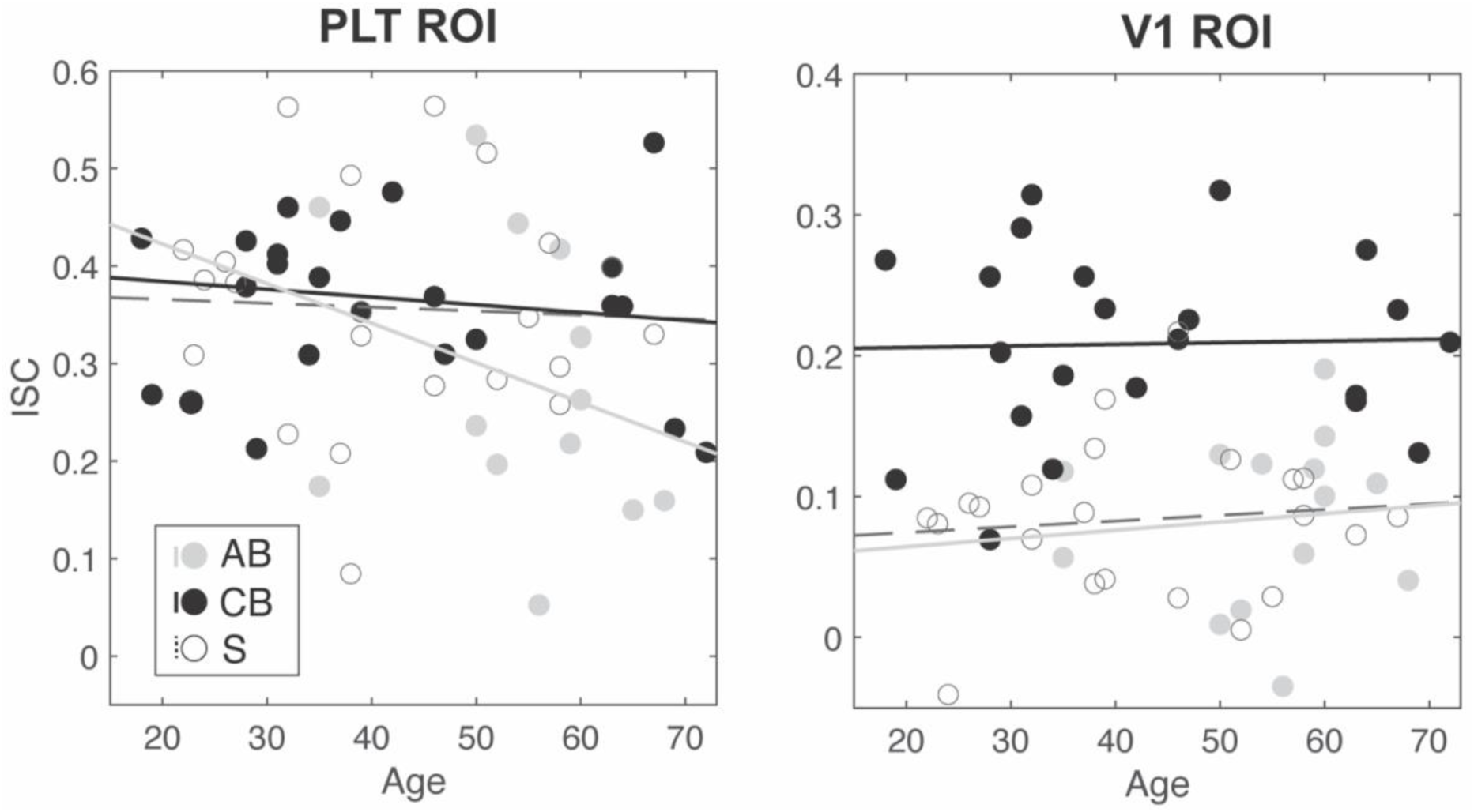
Relationship between participant age and ISC for the Audio-Movies condition in the PLT ROI (left) and the V1 ROI (right). Trend lines depict the linear relationship between age and ISC in each subject group. Participant age was not strongly correlated with ISC in either PLT (AB: *r*= −.28; CB: *r*= −.16; S: *r*= −.04) or V1 (AB: *r*= .09; CB: *r*= .03; S: *r*= 0.1).

In addition, in the V1 ROI, reliable ISC emerged for the narrative Pie-Man stimulus in the CB group (*r*=.20, *p*<.001) but not in the AB group (*r*=.001, *p*>.1) or the S group (*r*=.02, *p*>.1). The CB group ISC values for Pie-Man were reliably greater than both the AB group and the S group (both *p*<.001), while the AB and S groups did not differ from one another (*p*>.1). Additionally, V1 timecourses for the CB group showed greater synchrony for the intact Pie-Man stimulus versus Backwards Speech (*p*<.001). In contrast, ISC values for these stimuli did not vary for the S group (*p*>.1) or the AB group (*p*>.1). Group-by-condition interaction tests confirm that the difference in ISC values for Pie-Man versus Backwards Speech was greater in the CB group than in either the S group (*p*<.001) or the AB group (*p*<.001), while the S group and AB groups did not differ from one another (*p*>.1).

For the Sentence-Shuffle stimuli, V1 timecourses were synchronized for the CB group (*r*=.11, *p*<.001), but not for the AB group (*r*=-.004, *p*>.1) or for the S group (*r*=.01, *p*>.1). CB synchrony in response to the Sentence-Shuffle condition was greater than both AB synchrony (*p*<.001) and S synchrony (*p*<.001), while synchrony did not vary between the AB and S groups, (*p*<.01). Although ISC for the Sentence-Shuffle condition in the CB group was significant, V1 synchrony was greater for the Pie-Man condition (*p*<.001). In contrast, both the AB group and S groups showed no differences between ISC values for Pie-Man versus Sentence-Shuffle (both *p*>.1).

In the A1 ROI, there was significant inter-subject correlation (ISC) for the Backwards Speech condition in all three groups (S: *r*=.34; CB: *r*=.25; AB: *r*=.27, all *p*<.001), and ISC did not differ by group (all *p*>.1). In A1, Audio-Movies also showed significant ISC in all three subject groups, but ISC for the S group (*r*=.47) exceeded both the CB group (*r*=.37) and the AB group (*r*=.36, all *p*<.01), while the CB and AB groups did not differ from one another (*p*>.1). Tests for group-by-condition interactions revealed that the difference in ISC values for Audio-Movies versus Backwards Speech did not vary among subject groups (all *p*>.1).

For the PLT language ROI, both Backwards Speech and Audio-Movies induced reliable ISC in all groups (Backwards Speech: S: *r*=.13, CB: *r*=.13, AB: *r*=.14, all *p*<.001; Audio-Movies: S: *r*=.36, CB: *r*=.37, AB: *r*= .28, all *p*<.001). However, synchrony was higher for Audio-Movies than Backwards Speech in all groups (all *p*<.001). Synchrony for Backwards Speech did not vary by group (all *p*>.1). Synchrony for the Audio-Movies was significantly lower among AB participants than the CB group (*p*<.001), and marginally lower than the S group (*p*=.09). The difference in Audio-Movies synchrony for the CB versus S groups was not significant (*p*>.1). In group-by-condition interaction tests, the difference in ISC values for Audio-Movies versus Backwards Speech was reliably lower in the AB group than in the CB group (*p*=.03), and marginally lower than in the S group (*p*=.05).

Since the PLT ROI showed somewhat lower synchrony in the AB group, we performed a control analysis to determine whether reduced synchrony in V1 for the AB group relative to the CB group could be accounted for by globally lower synchrony. We performed a subset analysis using data from Movie 2 exclusively, for which ISC in PLT was matched across groups (all group pairwise comparisons *p*>.1). In the V1 ROI, AB ISC for Movie 2 was significantly lower than the CB group (*p*<.01) but did not differ from the S group (*p*>.1). This analysis shows that ISC in V1 is lower in AB relative to the CB group even for stimuli that are matched in PLT.

Neither the CB group nor the S group showed any reliable synchrony in any ROI during the resting state scan (all *p*>.1). Resting state data were only available for 10 of the 14 AB participants and in this subgroup the timecourses among AB participants during rest was negatively correlated in the A1 ROI (*r*=-.07, *p*<.01), the PLT ROI (*r*=-.07, *p*<.05), and the V1 ROI (*r*=-.09, *p*<.01). These results should be interpreted with caution, as only a relatively small subset of participants contributed resting state data to this analysis.

### V1 synchrony differences between CB and AB participants not accounted for by blindness duration or age

People who are born blind are on average blind for a longer time than people who become blind as adults. It is therefore possible that differences observed between CB and AB groups are related to blindness duration, rather than age of blindness onset. Contrary to this possibility, we found that among AB participants, there was no significant relationship between blindness duration and V1 ISC values (i.e., each AB participant to the rest-of-group average timecourse) (*r*=.-11, *p*>.1) or between blindness duration and between-group V1 synchrony with the congenitally blind group (i.e., each AB participant to the CB group-average timecourse) (*r*=-.03, *p*>.1) (Figure C2). This is despite the fact that duration varied considerably within the AB group (SD = 10.8, min= 6, max= 40).

Finally, we tested whether age could explain the relatively lower synchrony observed in the V1 of the AB group than in the other two groups, as AB participants were generally older than both the CB and S participants (see Methods). For this analysis, we ran linear regressions using age to predict Audio-Movies ISC values, and then tested whether the residual values, after accounting for the variance in ISC attributed to age, varied by group. After partialling out the effect of age, in the V1 ROI, the residual ISC values for the AB group remained significantly lower than CB after accounting for age (*p*< 0.0001) and did not differ from the S group (*p*> .1). By contrast, after partialling out age, PLT ISC values were marginally lower for AB versus CB (*p*= .10) but did not differ between the AB and S groups (*p*>.1). Age was not strongly correlated with ISC values for the AB group in either the V1 (*r*= −.09) or the PLT (*r*= −.28) ROI (Figure 3).

## Discussion

It was previously found that listening to meaningful naturalistic stimuli, including sound excerpts from popular live-action movies and a spoken narrative, but not low-level auditory stimuli (i.e., backwards speech), synchronizes activity in visual cortices across congenitally blind people. Synchrony observed in people who are congenitally blind is higher than among blindfolded sighted people (Loiotile et al., 2019; Van Ackeren et al., 2018). Here we report that in contrast to people who are born blind, people who become blind as adults show no more synchrony than blindfolded sighted adults, either for meaningful stimuli (movies and spoken narrative) or for backwards speech. By contrast, outside of the visual cortices, inter-subject correlations in the CB and S groups were largely similar, although AB individuals showed somewhat less synchrony in some higher-order areas, such as the lateral temporal cortex. Visual cortex synchrony in people who become blind as adults is not observed even after many years (e.g., four decades) of total blindness and does not change with blindness duration. No more synchrony is observed for an individual blind for forty years than for another blind for six. The absence of shared responses among people who become blind as adults suggests that the human visual cortex has a special capacity to assume novel functions during a sensitive period of development.

### Evidence for a sensitive period in visual cortex plasticity during naturalistic listening

The reduced inter-subject correlations in visual cortex among the AB group, relative to the CB group, observed in the current study is consistent with previous experimental studies reporting reduced or absent cross-modal activity in adult-onset blind individuals (Bedny et al., 2010, 2012; Cohen et al., 1999; Jiang et al., 2016; Kanjlia et al., 2019; Pant et al., 2020; Sadato et al., 2002). We find that both in whole-cortex analyses and in focused analyses examining V1 responses, the AB group was different from the CB and no different from the sighted. The current results with naturalistic auditory stimuli further suggest that previously observed differences between AB and CB groups are not idiosyncratic to a particular task or cognitive process. A key property of these naturalistic stimuli is that they are rich with information across different cognitive levels, ranging from low-level perceptual features (e.g., changes in pitch and volume) to intermediate linguistic properties (e.g., individual words and sentences) to higher-order semantic content (e.g., narrative plot and character intent). None of these features appear to induce synchrony in the AB population. We also found no evidence that visual cortex synchrony in AB (or CB) individuals is driven by low-level auditory features (i.e., backwards speech). Together, these results suggest that responses to auditory stimuli in adult-onset blind individuals are weaker and/or inconsistent across individuals, relative to people born blind.

Both the adult-onset blind and blindfolded sighted groups showed reduced synchrony relative to the CB participants, to a similar degree. Nevertheless, region of interest analyses in V1 revealed an equally low but significant degree of synchrony among blindfolded sighted and adult-onset blind people. This observation is consistent with previously observed responses to nonvisual information in visual cortices of sighted individuals. During visual tasks in sighted people, responses in the occipital lobes are modulated by top-down task demands (Gazzaley et al., 2007; Ruff et al., 2006; Serences & Yantis, 2007; Waskom et al., 2014). Even in the absence of visual input, the visual cortices are recruited when sighted participants engage in mental imagery (Hindy et al., 2015; Hsu et al., 2012; Stokes et al., 2009), and when visual stimuli are anticipated or remembered but not currently present (Kastner et al., 1999; Sergent et al., 2011). Nonvisual stimuli, such as tactile and auditory input, have also been shown to elicit visual cortex activity in sighted people (Facchini & Aglioti, 2003; James et al., 2002; Merabet et al., 2004; Merabet et al., 2008; Sathian, 2005; Seydell-Greenwald et al., 2020; Vetter et al., 2014; Voss et al., 2016; Zangaladze et al., 1999). These findings demonstrate that nonvisual input can activate the visual cortices independent of one’s vision status.

The behavioral and cognitive significance of cross-modal responses in sighted and adult-onset blind individuals is currently unknown; it is possible that such activity is epiphenomenal with respect to behavior in these groups and gains behavioral significance in people who are born blind. At least one TMS study comparing adult-onset and congenitally blind adults supports this view (Cohen et al., 1999; see also Amedi et al., 2004). The present results, together with prior literature further show that cross modal responses in people born blind are more anatomically extensive and robust, occur under different stimulus conditions, and are more sensitive to subtle cognitive manipulations (e.g., Pant et al., 2020; Kanjlia et al., 2016; Lane et al., 2015). Thus, while cross-modal effects are present in visual cortices of all people, early blindness substantially modifies the character of these responses. Nevertheless, the existence of cross-modal responses in both blind and sighted adults suggests that in both populations, there are pathways that transmit nonvisual information to the visual cortices. Plasticity observed in people who are congenitally blind likely occurs by functionally modifying the anatomical mechanisms present in sighted and blind people alike (Collignon et al., 2011; Merabet et al., 2004; Pascual-Leone & Hamilton, 2001; Sathian & Stilla, 2010; Stilla et al., 2008; Wolbers et al., 2011).

### Caveats: Some adult-onset cross-modal responses not ruled out by naturalistic designs

The present findings provide compelling evidence that adult-onset blindness does not enhance shared visual cortex responses for naturalistic auditory stimuli. However, we cannot definitively rule out the possibility that *some* cognitive process would reveal cross-modal responses in people who become blind as adults that are different from blindfolded sighted people. Although the space of possible response properties sampled by naturalistic auditory stimuli is relatively wide, it is by no means exhaustive. One kind of manipulation that is lacking in the current study is an explicit task with overt responses. Naturalistic listening is a relatively passive and effortless task that does not require making any overt responses during stimulus presentation. Synchrony observed in naturalistic paradigms is unlikely to be driven by task demands. However, if visual cortices of adult-onset blind people become active during decision making, or motor function, then a naturalistic listening paradigm would fail to induce such activity. Indeed, several studies have found some activity in visual cortices of adult-blind onset individuals during effortful tasks that involve making on-line decisions. For example, cross-modal activity has been reported when adult-onset blind participants make button presses or verbal responses to indicate semantic judgments in response to individual words and sentences (Aguirre et al., 2016; Burton et al., 2003; Burton, Snyder, Diamond, et al., 2002; Burton & McLaren, 2006; Pant et al., 2020). Notably, even when observed, such responses are weaker than those seen in visual cortices of people born blind during the same tasks. Nevertheless, understanding the role of decision making and motor output in driving cross-modal activity in visual cortex of adult-onset blind and blindfolded sighted people merits further research.

A further element of the present design worth considering is that lack of synchrony among AB participants in our study could reflect individual variability in this population, rather than a lack of cross-modal responses in each individual. Unlike task-based designs, synchrony paradigms, such as this one, rely on activity occurring in relatively similar regions across people. Thus, if entirely different parts of visual cortex are responsive in different adult-onset blind individuals, synchrony would not be observed in the group. The current findings could thus indicate either that there is a sensitive period for developing cross-modal responses to naturalistic sounds or instead that there is a sensitive period for having anatomical systematicity of such responses across individuals. One way to distinguish between these possibilities in future studies would be to look for correlated responses to repeated listening of naturalistic auditory stimuli within each AB individual.

### Summary and conclusions

In conclusion, visual cortices of people born blind show robust synchrony for meaningful naturalistic auditory stimuli. Here we report that vision during development severely attenuates this shared activity. “Visual” cortices of individuals who were sighted during childhood, lost vision as adults and have, in some cases, been blind for decades, remains functionally similar to the visual cortices of people who are sighted. These results support the view that the cognitive flexibility and systematic reorganization of visual cortices is limited by sensitive periods during development.

## Acknowledgments

This work was supported by the National Institute of Health (NEI, R01 EY027352-01 to M.B.) and a National Science Foundation Postdoctoral Research Fellowship (SBE SPRF 1911650 to E.M.). We thank the blind and sighted individuals who participated in this study and we are grateful toward the blind community for its support of this research. We also thank the F. M. Kirby Research Center for Functional Brain Imaging at the Kennedy Krieger Institute.

## Appendix A

**Table A1.** Demographic information for adult-onset participants.

| Participant | Gender | Age | Blindness cause | Light perception | Education | Age functional loss began | Age reached current vision level | Blindness duration at current level |
| --- | --- | --- | --- | --- | --- | --- | --- | --- |
| AB_01 | M | 50 | Optic nerve neuropathy | None | BA | 21 | 34 | 16 |
| AB_02 | F | 65 | Autoimmune disorder | None | AA | 39 | 57 | 8 |
| AB_03 | M | 52 | Glaucoma | None | HS | 38 | 38 | 14 |
| AB_04 | M | 56 | RP | Minimal | BA | 33 | 35 | 21 |
| AB_05 | M | 50 | Trauma | None | JD | 19 | 19 | 31 |
| AB_06 | F | 35 | RP | Minimal | MA | 17 | 25 | 10 |
| AB_07 | F | 68 | Glaucoma | Minimal | BA | 48 | 49 | 19 |
| AB_08 | M | 59 | Trauma | None | BA | 50 | 50 | 9 |
| AB_09 | F | 60 | Diabetic retinopathy | None | MA | 19 | 20 | 40 |
| AB_10 | F | 60 | Diabetic retinopathy | None | BA | 50 | 52 | 8 |
| AB_11 | F | 58 | RP | None | SC | 35 | 54 | 4 |
| AB_12 | F | 35 | Stevens-Johnson Syndrome | None | BA | 25 | 25 | 10 |
| AB_13 | F | 54 | Glaucoma | None | DPT | 17 | 24 | 30 |
| AB_14 | F | 60 | Uveitis | None | MA | 48 | 50 | 10 |

RP = retinosis pigmentosa; BA = Bachelor of Arts; AA = Associates Degree; JD = Juris Doctorate; SC = some college; DPT = doctorate of physical therapy

**Table A2.** Behavioral performance on the post-scan multiple-choice questionnaire, separated by subject group and stimulus condition

| Subject Group | Pie-Man | Movie 1 | Movie 2 | Movie 3 |
| --- | --- | --- | --- | --- |
| S (n=22) | 4.67 (0.73) | 3.81 (1.44) | 4.10 (0.89) | 3.52 (1.54) |
| CB (n=22) | 4.77 (0.53) | 4.00 (1.31) | 3.95 (1.65) | 3.64 (1.14) |
| CB subset (n=14) | 4.64 (0.63) | 3.42 (1.34) | 3.35 (1.82) | 2.93 (0.73) |
| AB (n=14) | 4.07 (1.27) | 3.36 (1.50) | 3.43 (1.22) | 3.00 (1.30) |

Scores for each stimulus condition is out of a total of five points. Mean scores are listed with standard deviations in parentheses. In post-hoc t-tests for each condition, behavioral performance did not differ between subject groups, all $ps > 0.05$ . S = sighted; CB = congenitally blind; AB = adult-onset blind

## Appendix B

**Figure B1.**
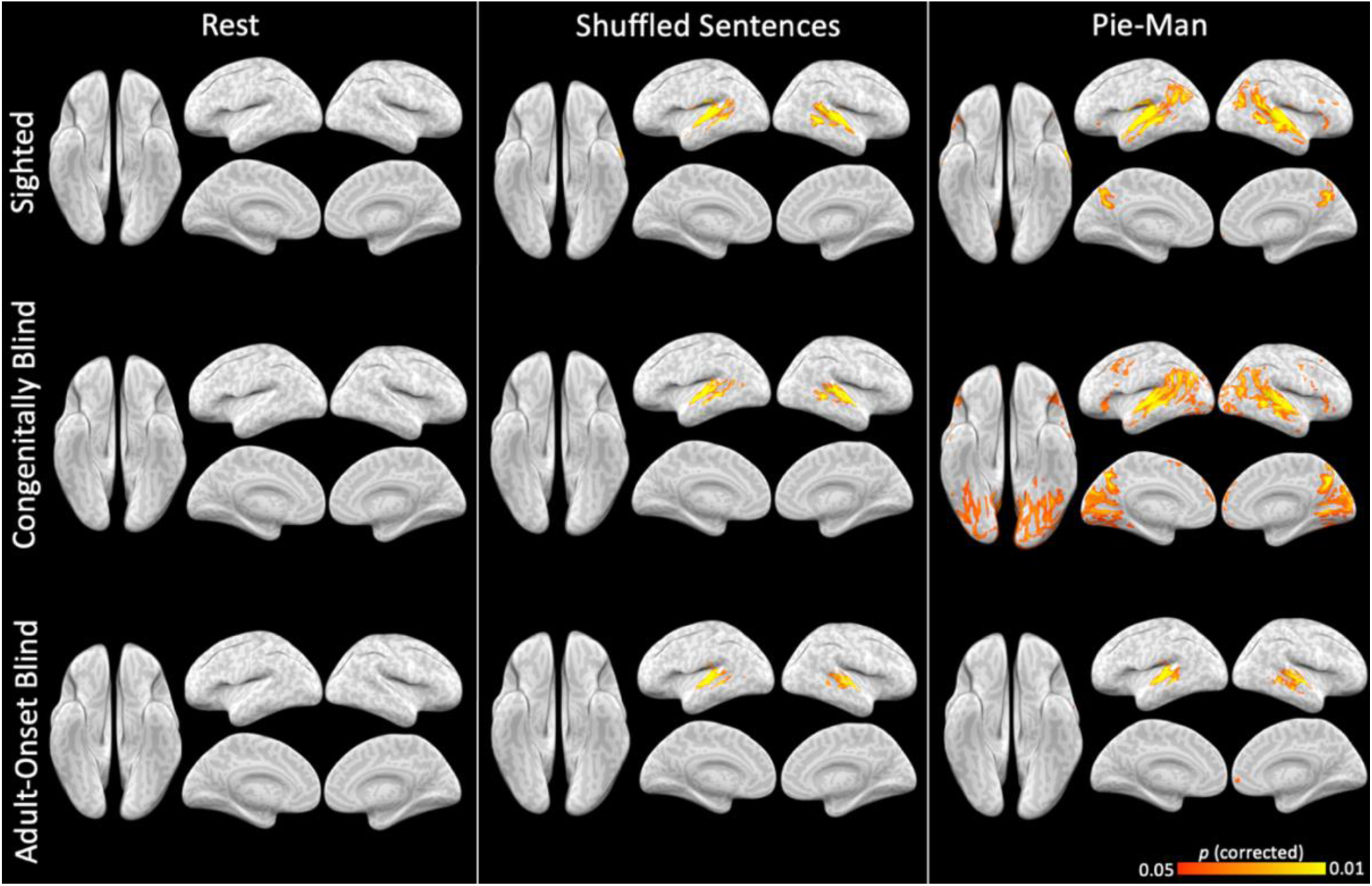
Whole-brain inter-subject correlation (ISC) maps for the resting state scan, the shuffled sentences version of the Pie-Man comedy sketch stimulus, and the intact version of the Pie-Man stimulus in each subject group. Voxel-level synchronization values are corrected for multiple comparisons, *p*<.05.

**Figure B2.**
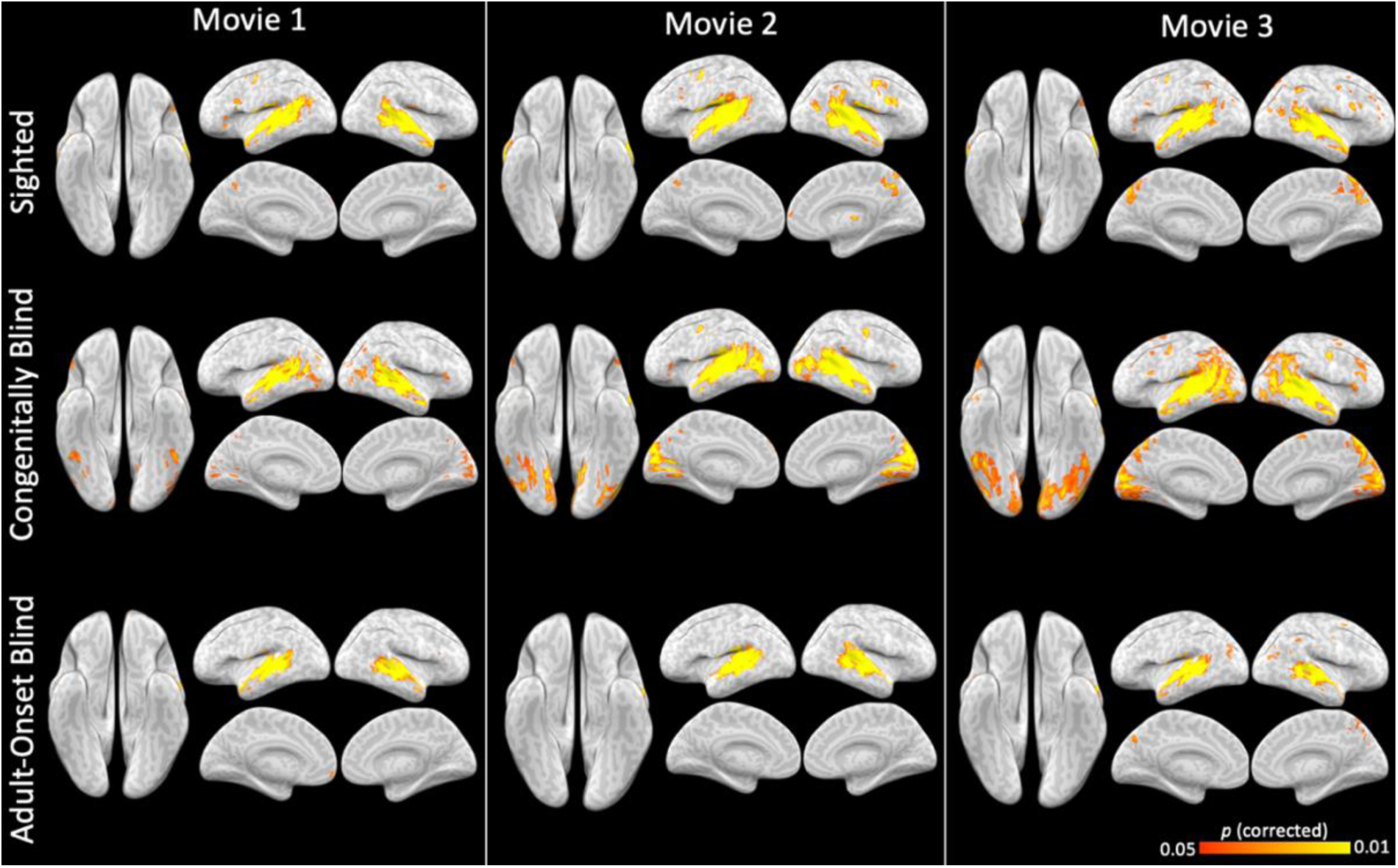
Whole-brain, voxel-level ISC maps elicited by each individual movie condition in each vision group, corrected for multiple comparisons at *p*<.05. Movie 1 = “Blow Out”; Movie 2 = “The Conjuring”; Movie 3 = “Taken”

## Appendix C

**Figure C1.**
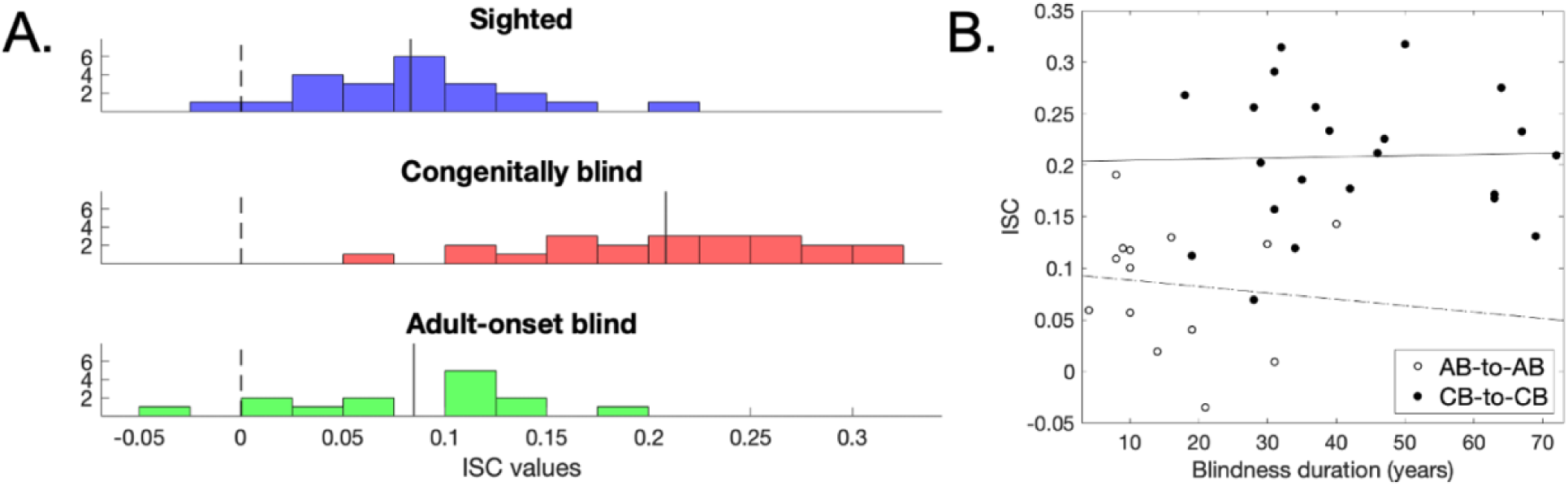
Participant-level ISC values in the V1 ROI for the Audio-Movies condition. A: Distribution of ISC values across individuals in each subject group. The solid vertical line on each group’s histogram corresponds to the group-average ISC value and the dashed vertical line shows the location of an ISC value of zero. B: Relationship between duration of blindness (in years) and inter-subject correlation (ISC) in the V1 ROI during the Audio-Movies stimuli. Each datapoint represents one individual. Linear trends are depicted for each group (AB: *r*= −.11, dashed line; CB, *r*= .03, solid line). For the CB group, blindness duration is equivalent to their age at time of test.

**Figure C2.**
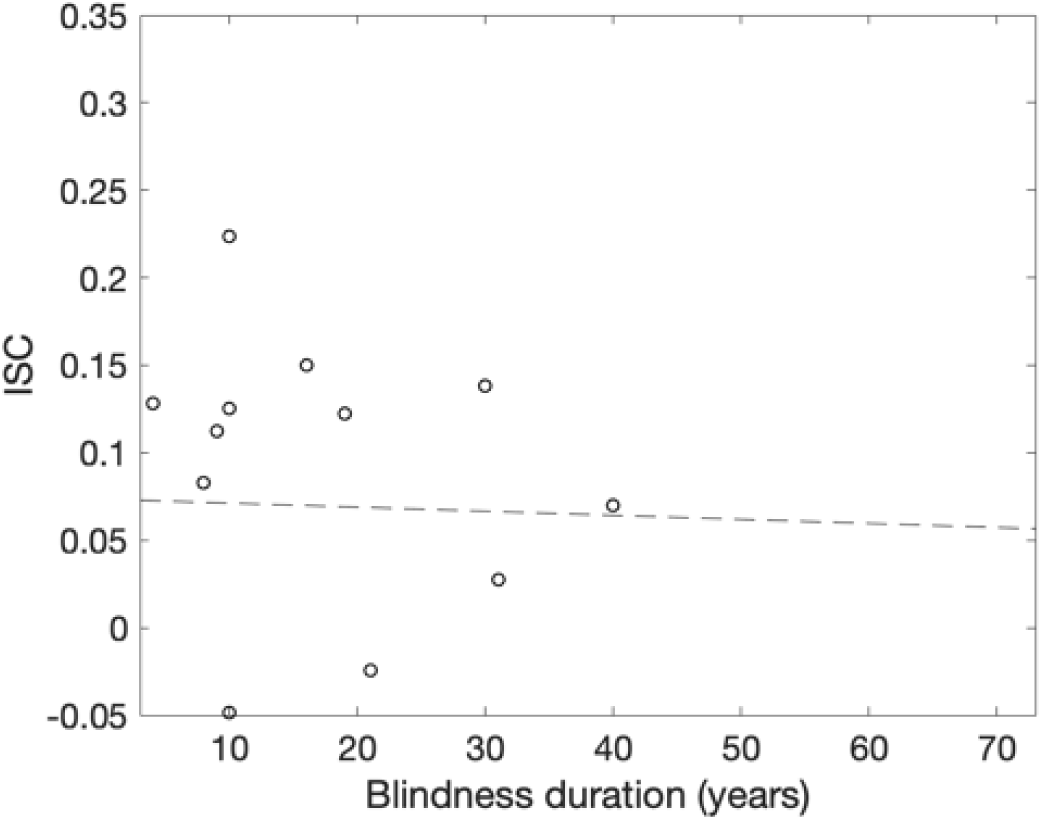
Relationship between each AB participant’s duration of blindness and their between-group ISC value (AB-to-CB) in the V1 ROI during the Audio-Movies stimuli. Here, the ISC value reflects the correlation between each individual AB participant’s timecourse and the group-average timecourse for the CB group. Each datapoint represents one AB individual. Across AB participants, duration of blindness was not correlated with synchrony to the CB group in V1, *r*= −.03).

